# Distinct Microstructural Heterogeneities Underpin Specific Micromechanical Properties in Human ACL Femoral and Tibial Entheses

**DOI:** 10.1101/2023.08.16.553628

**Authors:** Jinghua Fang, Xiaozhao Wang, Huinan Lai, Wenyue Li, Zongyou Pan, Renwei Mao, Yiyang Yan, Chang Xie, Junxin Lin, Wei Sun, Rui Li, Jiajie Wang, Jiacheng Dai, Kaiwang Xu, Xinning Yu, Tengjing Xu, Wangping Duan, Jin Qian, Hongwei Ouyang, Xuesong Dai

## Abstract

The anterior cruciate ligament (ACL) is anchored to the femur and tibia by a specialized interface tissue called the enthesis, which transfers forces in multiple directions and magnitudes without accruing fatigue damage during loading cycles over a lifetime. However, the precise structural and mechanical characteristics of the ACL femoral enthesis (FE) and tibial enthesis (TE) and their intricate interplay are unknown. In this study, we identified two ultrathin-graded mineralization regions in the FE (∼21 μm) and TE (∼14 μm), both of which exhibited distinct biomolecular compositions and mineral assembly patterns. FE interface exhibited progressively maturing hydroxyapatites (HAps), whereas minerals at the TE interface region changed from an amorphous phase (ACP) to HAps with increasing crystallinity. The LC-MS/MS results revealed that MGP protein uniquely enriched at the TE interface may be favorable for stabilizing ACP, while CLEC11A enriched at the FE interface could facilitate osteogenesis of the interface. The finite element analysis results indicated that the FE model was more resistant to shearing, while the TE model facilitated tensile resistance. It suggested that the great discrepancy in biomolecular expression and the corresponding mineral assembling heterogeneities together contributed to the superior mechanical properties of both the FE and TE models. These findings provide new perspectives regarding the management of ACL injury and the development of high-performance interface materials.

## INTRODUCTION

The enthesis, a component facilitating musculoskeletal motion, is a complex structure that acts as a connection between soft and hard tissues.^1,2^ The anterior cruciate ligament (ACL) serves as a vital mechanical stabilizer within the knee joint, linking the femur to the tibia via the ACL-femoral enthesis (TE) and ACL-tibial enthesis (TE).^3^ ACL injuries are prevalent, and only 16% occur at FE and 1% at TE.^4,5^ Given the limited capacity for natural ACL healing, surgical intervention is necessary. However, the intricate reconstruction of the ACL-bone interface presents challenges, affecting the graft’s durability and knee function over time.^6,7^ Despite the potential risk of stress concentration and subsequent fracture at the interface between soft and hard materials,^8^ the FE and TE demonstrate remarkable toughness and resilience against combined tension, shear, and torsional forces, with multidirectional stresses applied to the FE and relatively unidirectional stress on the TE.^1,9^ This resilience is attributed to the multi-scale toughening mechanism of extracellular matrix (ECM) present in natural materials.^10–13^ Thus, the analysis of the interplay between organic and inorganic structural components at the ACL-bone interface offers valuable insights into its high-strength mechanical conduction mechanisms.

Recent studies have focused extensively on the structural components of the ECM at the tendon/ligament-bone interface.^2,11,14–18^ This interface exhibits a linear transition from tendon/ligament to fibrocartilage and then to subchondral bone (SB).^19,20^ The fibrocartilage zone is further divided into nonmineralized (NFC) and mineralized (MFC) regions.^21^ The gradual increment in mineral contents across the gradient mineralized structure is crucial, as it results in a substantial increase in tissue modulus without causing stress concentration.^13,22–25^ Despite the utilization of various micro-nano analytical techniques to investigate the nanoscale structure and composition of bone, ligament, and tendon,^26–29^ there is still limited knowledge regarding the multi-scale structure assembly of the gradient mineralized interface.^30,31^ Further research is needed to clarify the multiscale composition and assembly of the graded minerals, as well as the precise biomolecular composition and stiffness transition of interface tissue. These findings will lay the groundwork for designing soft-to-hard interfaces and determining parameters for current ACL-bone regeneration and repair techniques.^32,33^

Consequently, this study employs a variety of advanced techniques (Figure S1), including scanning electron microscopy (SEM), high-angle annular dark field scanning transmission electron microscopy (HAADF-STEM), focused ion beam-SEM (FIB-SEM), Raman spectroscopy, high-resolution TEM-electron energy loss spectroscopy (HRTEM-EELS), and nanoindentation, along with liquid chromatography-tandem mass spectrometry (LC-MS/MS), to examine the location-specific graded mineral composition and assembly of FE and TE and analyze their respective effects on the overall mechanical functions in human knee joints.

## RESULTS AND DISCUSSION

### Structural Transition and Mechanical Performance of FE and TE Interfaces

Normal FE and TE tissues were collected (Figures 1A and S2A) and identified using histological staining (Figures 1C and S3). Using high-resolution microcomputed tomography (miro-CT) and SEM, we then studied the microstructures and visualized the macro interlocking structures of interfacial tissues of FE and TE. Density-dependent color electron micrographs (DDC-SEM) depicted the microstructural alterations at the interface between NFC regions (green) and highly mineralized regions (red) (Figures 1B, D, S2B, and S4).^12,34^ In lieu of a straight line, the interface in both FE and TE samples displays a wave-like curvature,^35,36^ with the femoral side exhibiting a wider graded mineralization interface.^1,37^ We identified specific fractional mineralized regions of FE and TE interfaces by employing energy dispersive X-ray (EDX) line scanning and surface scanning (Figures 1E, F, and S5). The exponential increment curve of Ca and P content in the FE interface region yielded an ultra-thin region of 21.11±2.59 μm and 14.56±3.19 μm in the wave crest and trough region, respectively. Ultrathin regions of 13.84±3.41 μm and 6.77±1.70 μm were obtained in the crest and trough regions of the TE interface, respectively, demonstrating the seamless transition from NFC to highly mineralized regions at both interfaces.^34^

**Figure 1.**
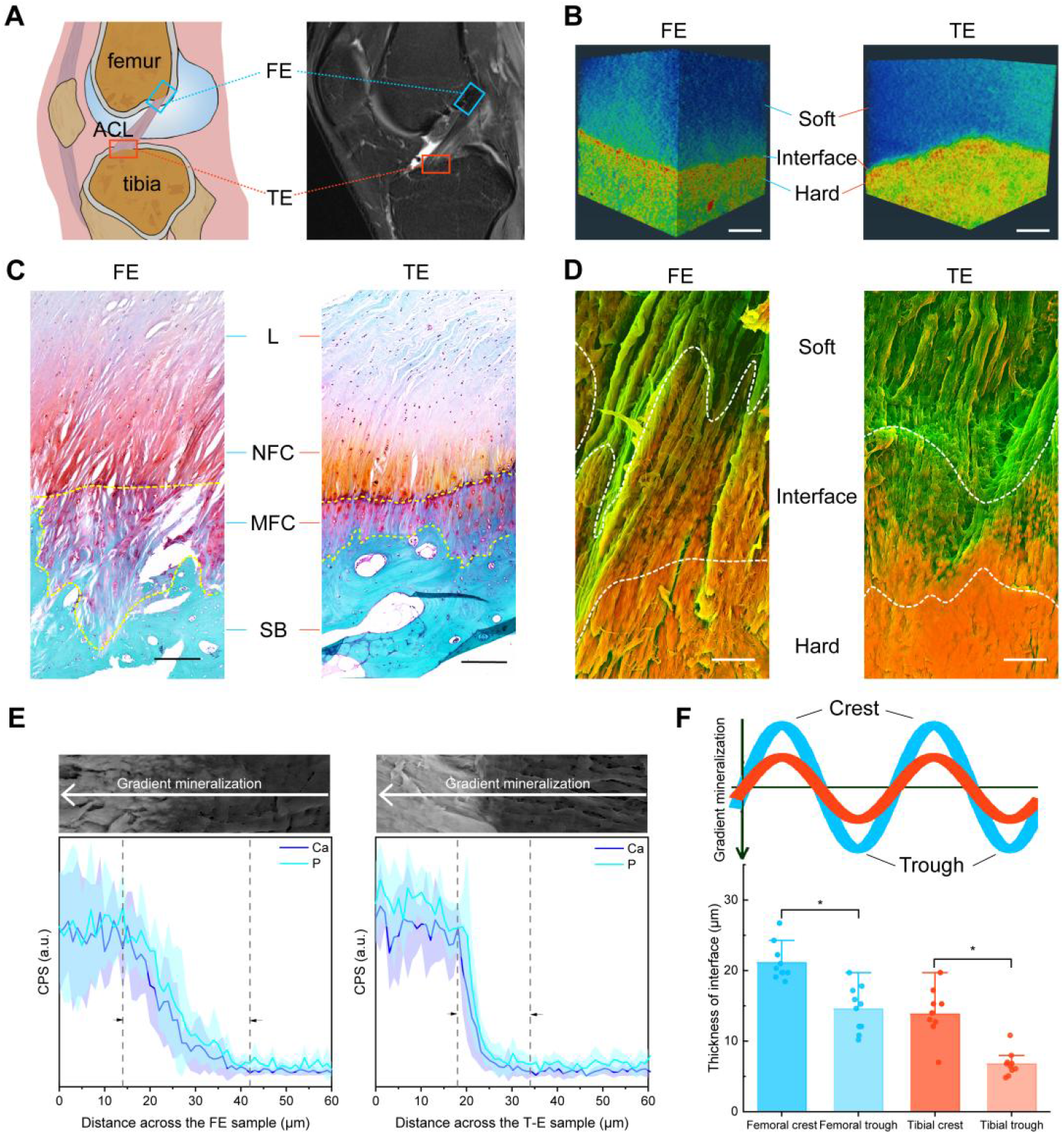
Structural transition of FE and TE Interfaces. (A) Schematic and magnetic resonance imaging (MRI) illustration of human knee showing femur-ACL-tibia complex. (B) Reconstructed 3D volumetric representation of the structures including ligament, ACL-bone interface and SB tissues. The micrographs were colored in postprocessing by Amira 2020 (the original images, and processing details see Figure S2B). (C) Safranin O/Fast Green (SO) staining images showing histological features of different samples (FE and TE). (D) DDC-SEM micrographs of FE and TE interface tissues presenting gradual mineral distributions with distinct morphologies. The micrographs were colored in postprocessing by combining images obtained from backscatter electron and secondary detectors (red-minerals, green-organic materials, the original images, and processing details see Figure S4). (E) SEM images and the corresponding EDX line scan (Ca and P) (down) identifying FE and TE interface. (F) The thickness of the two mineralized interfaces quantified by EDX line scan in (D). Scale bar in (B), 30 μm; (C), 200 μm; (D), 5 μm.

Significantly, the rough mineralized interface potentially contributes to enhanced toughness at the tendon-to-bone insertion site,^35^ and the thickness of the interface plays a critical role in determining mechanical properties.^38^ Therefore, we conducted an exhaustive evaluation of the mechanical performance of two interface tissues, FE and TE. The stress-strain curves clearly illustrated the distinct mechanical properties of these samples. Moreover, no fissures were observed in the region of entheses (Figure S6). The nanoindentation test results indicated a substantial increase in tissue modulus at the interface for both FE and TE samples.^39,40^ The tissue modulus of the FE sample increased from 39.72 ± 4.74 MPa to 760.39 ± 39.90 MPa over a 20 μm thickness interface, whereas the tissue modulus of the TE sample increased exponentially from 27.19 ± 4.53 MPa to 1261.98 ± 217.77 MPa over a 10 μm thickness interface (Figure 2A-C). To further explore the nanoscale mechanical properties, we employed atomic force microscopy (AFM) to examine the low (i), medium (ii), and highly mineralized (iii) regions of FE and TE samples (Figure 2D-F).^41^ The distribution of tissue hardness in region i of the FE samples exhibited greater variation than that of the corresponding regions in the TE samples. Notably, the medium mineralized region (zone ii) of the TE sample displayed a distinctive hardness peak of 1.82 GPa, which can be ascribed to the narrower mineralization gradient region in TE and comprised zone iii within the scanned area. These distinct hardness distributions in each mineralized zone are consistent with the microstructures depicted in Figure 1.

**Figure 2.**
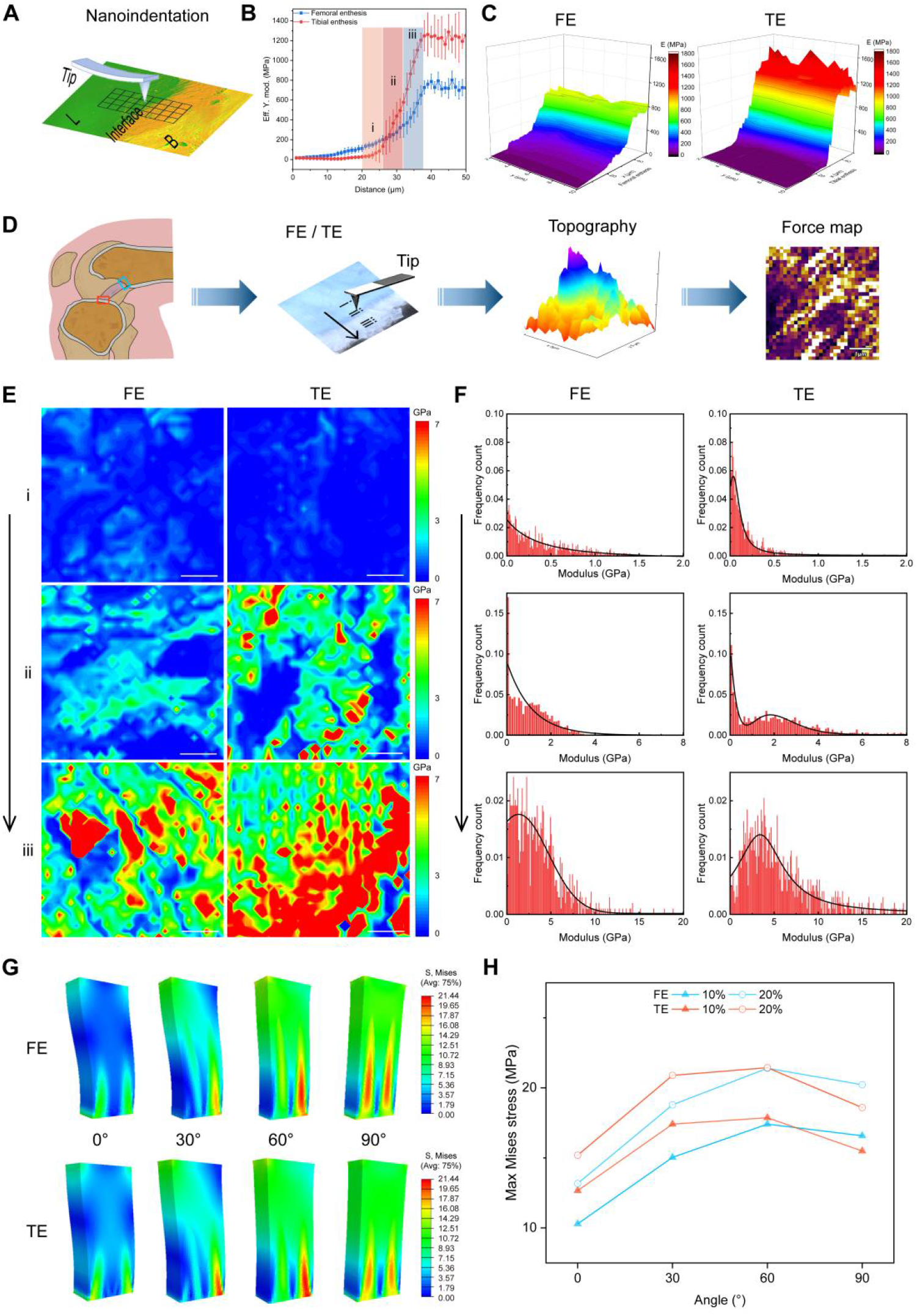
Correlating the local mechanical response of FE and TE. (A) Schematic view of nanoindentation test for FE and TE. (B) Average efficient modulus of FE and TE tissues as a function of distance from ligament. (C) Efficient modulus maps of the rectangular area in (A) (resolution: 1 μm in x-axis and 2 μm in y-axis). (D) Schematic diagram showing the test of nanomechanical response interface regions from less mineralized region (i) to intermediate mineralized region (ii), and to the highly mineralized region (iii) using AFM (0.16×0.16 μm step). (E) Consecutive AFM stiffness maps of selected regions in D (i,ii and iii) at interface tissues for both samples. (F) Corresponding stiffness distributions across calcified cartilage tissues for different samples. (G) Stress distribution under tension conditions in different directions. (H) The max von Mises stress on FE and TE under tension conditions in different directions. Scale bar in (E), 1 μm.

In order to meet the specific mechanical requirements at each insertion site, the microscopical and macroscopical structures of the entheses vary based on their anatomical location.^42,43^ Previous studies have shown that FE had a larger MFC tissue area and a more acute ligament attachment angle than TE, with FE being subjected to multidirectional stresses and TE to relatively unidirectional ones.^1,9,37^ Finite element analysis (FEA) was used to further validate the mechanical superiority of both models based on the factors of insertion depth, structure, and mechanical gradient alteration in FE and TE (Figure S7).^44^ The FE model exhibited enhanced shear resistance, whereas the TE model exhibited greater tensile resistance. Specifically, in the force direction of 0°, the maximum force value of FE was lower than that of TE, whereas in the force direction of 90°, the maximum force value of FE exceeded that of TE (Figure 2G, H and movie S1).

### Multiscale Mineral Transformations at FE and TE Interfaces

Previous studies have demonstrated that spatial gradients in mineral distribution have the potential to reduce stress concentration during joint movement.^24,45,46^ This prompted us to examine the multiscale mineral transformations at the interfaces of FE and TE using HAADF-STEM and FIB-SEM (Figure 3). Both FE and TE samples exhibited graded areas of mineralization, with the FE sample showing noticeably denser mineralization (Figure 3A and B), a finding supported by SEM, EDX-line scan and TEM images (Figures 1D, E and S8). HAADF-STEM and FIB-SEM images of the FE sample revealed three distinct mineral morphologies: loosely fibrous particles, well-defined particles that grew in size and fused into platelets, and finally formed a compact accumulation. At the interface, the TE sample exhibited three typical mineral forms: sparsely distributed small spheroids that progressively fused into a bulk structure, resulting in dense packing (Figure 3C, D, E, and F). In the mineralization front region (zone i and ii), the minerals in the FE sample had a larger aspect ratio and denser distribution while those in the TE sample had an aspect ratio closer to 1 (Figure S9). Several studies have demonstrated that the addition of micro- or nano-particles can effectively improve the stiffness or Young’s modulus.^47,48^ In addition, experimental and atomic simulation studies have demonstrated that the elastic properties of nanoscale materials depend on their size.^49–51^ The graded distribution of minerals at the interface improves the adhesion of dissimilar tissues,^52^ and the minerals in the FE sample with larger aspect ratios may exhibit an increase in tensile adhesive energy,^53^ thereby effectively anchoring the ACL to the femur under multidirectional stresses.

**Figure 3.**
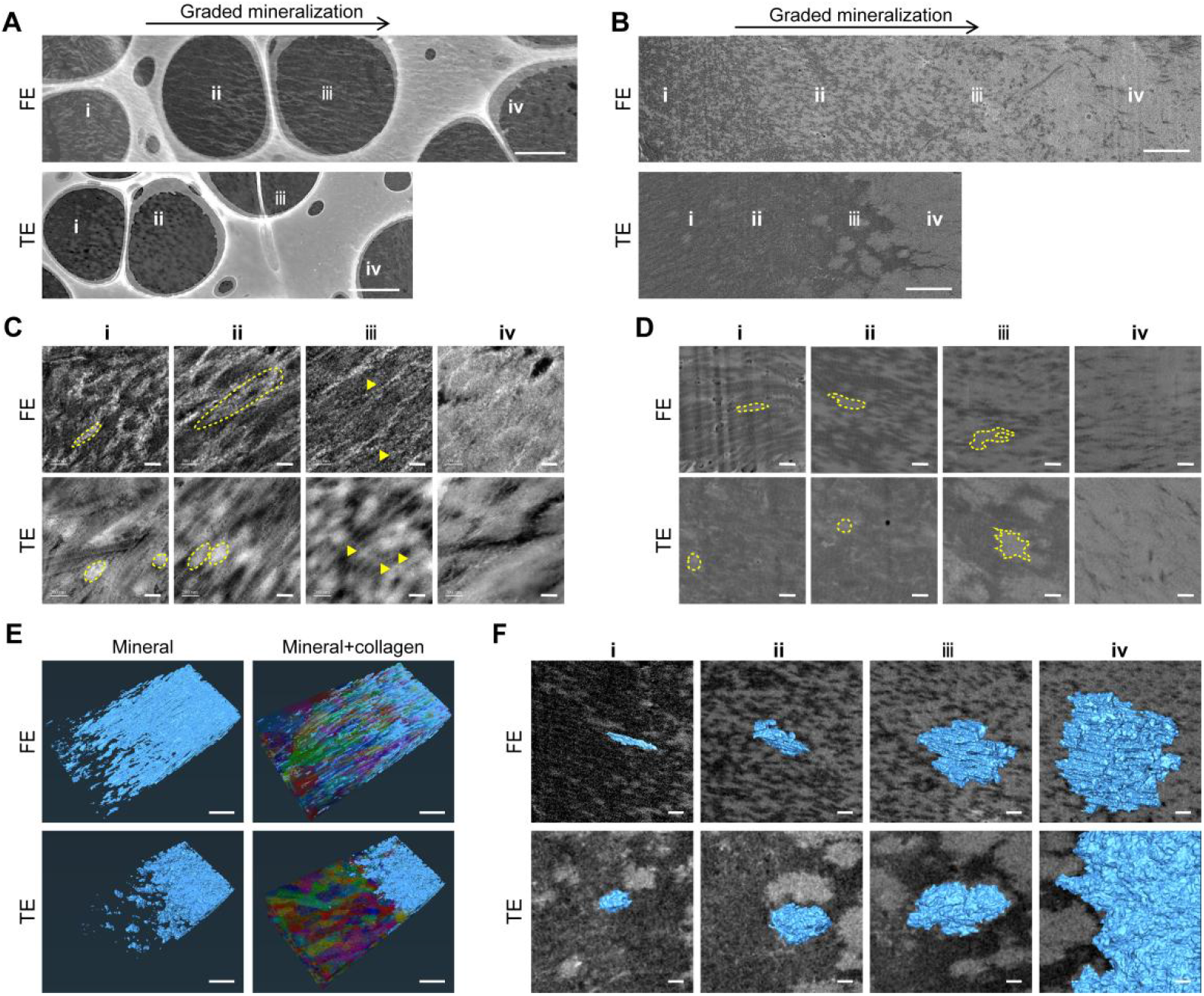
Multiscale structures of mineral transformations in FE and TE samples. (A) HAADF-STEM images showing the varied distributions and morphologies of minerals across both interfaces. (B) FIB-SEM images depicting the progression of morphologies, distributions and assemblies of mineral particles at FE and TE interfaces. (C) Corresponding enlarged STEM images in different regions in (A) present the transition of mineral morphologies and distributions at both interfaces. (D) Corresponding enlarged BSE images at different sites in (B) showing multiple minerals with various morphologies dispersed in the matrix. (E) 3D rendering of the mineralization front showing distinct mineral assembling patterns for FE and TE. (F) 3D rendering of typical mineral particles intersecting with FIB-SEM image planes at the mineralization front of the interface region, demonstrating the mineral appearance and aspect ratio in different samples. Scale bar in (A, B), 2 μm; (E), 1 μm; (C, D, F), 200 nm.

### Crystallographic Features and Chemical Analysis of Minerals at the FE and TE Interfaces

Apart from the gradient structures, the crucial material motif of the interface tissue also lies in the grading of local components.^24^ The diverse mineral morphologies observed in FE and TE interfaces necessitated a comprehensive compositional analysis. Thus, in this study, we conducted a high-resolution examination of mineral composition and spatial distribution at both the microscale and nanoscale in FE and TE. Raman spectroscopy was employed to assess the mineral phase at the FE and TE interfaces, which revealed the presence of carbonated hydroxyapatite (HAp) with PO_4_ ^3-^ v1 symmetric stretching at 960 cm^−1^ as well as a weak planar vibration of CO_3_ ^2-^ v1 peak at 1071 cm^−1^.^54,55^ Interestingly, a contralateral stretch at 957 cm^−1^ was observed in the mineral located in the front region of the TE interface (Figure 4A). This phenomenon can be attributed to the coexistence of HAp precursors and HAp, as further validated by stimulated Raman scattering microscopy (SRS). SRS spectra exhibited a peak at 950 cm^−1^ (ACP) in the initial region of the TE interface (Figure S10), and corresponding SRS imaging demonstrated the distribution of ACP and HAP at the TE interface, whereas no ACP was observed at the FE interface (Figure 4B).^54–56^ Considering that the configuration and position of PO_4_ ^3−^ at 960 cm^−1^ as well as CO_3_ ^2−^ band at 1071 cm^−1^ provide significant insights into mineral crystallinity, component proportions, and ionic substitutions, we generated composition maps of HAp and matrix to better comprehend their spatial distribution across both interfaces. These maps showed a gradual increase in HAp content and alternate carbonate content, demonstrating wider regions of graded MFC at the FE interface compared to the TE interface (Figure S11A and B). However, a reversed trend was observed in the CO_3_ ^2−^/PO_4_ ^3−^ ratio (Figure S11C). Further analysis of the interface divided it into four distinct regions (Figures 3A, C and S8), wherein a meticulous elemental analysis of the minerals in each region was conducted (Figure S12). Our findings indicated a significant increment in calcium, phosphorus, and oxygen content, accompanied by a decrease in carbon content, indicating an enhanced level of mineralization.^57^ Overall, the results of this study revealed a depth-dependent distribution of carbonate-substituted HAp at both the FE and TE interfaces, with a wider interface observed in FE samples compared to TE samples and a transformation of ACP to HAp in TE samples. The presence of amorphous calcium phosphate contributes to improved crack resistance,^56^ which may explain the higher structural integrity of TE. These findings were noteworthy as they highlight micro-scale spatial gradations in mineral composition and crystallinity at the FE and TE interfaces.

**Figure 4.**
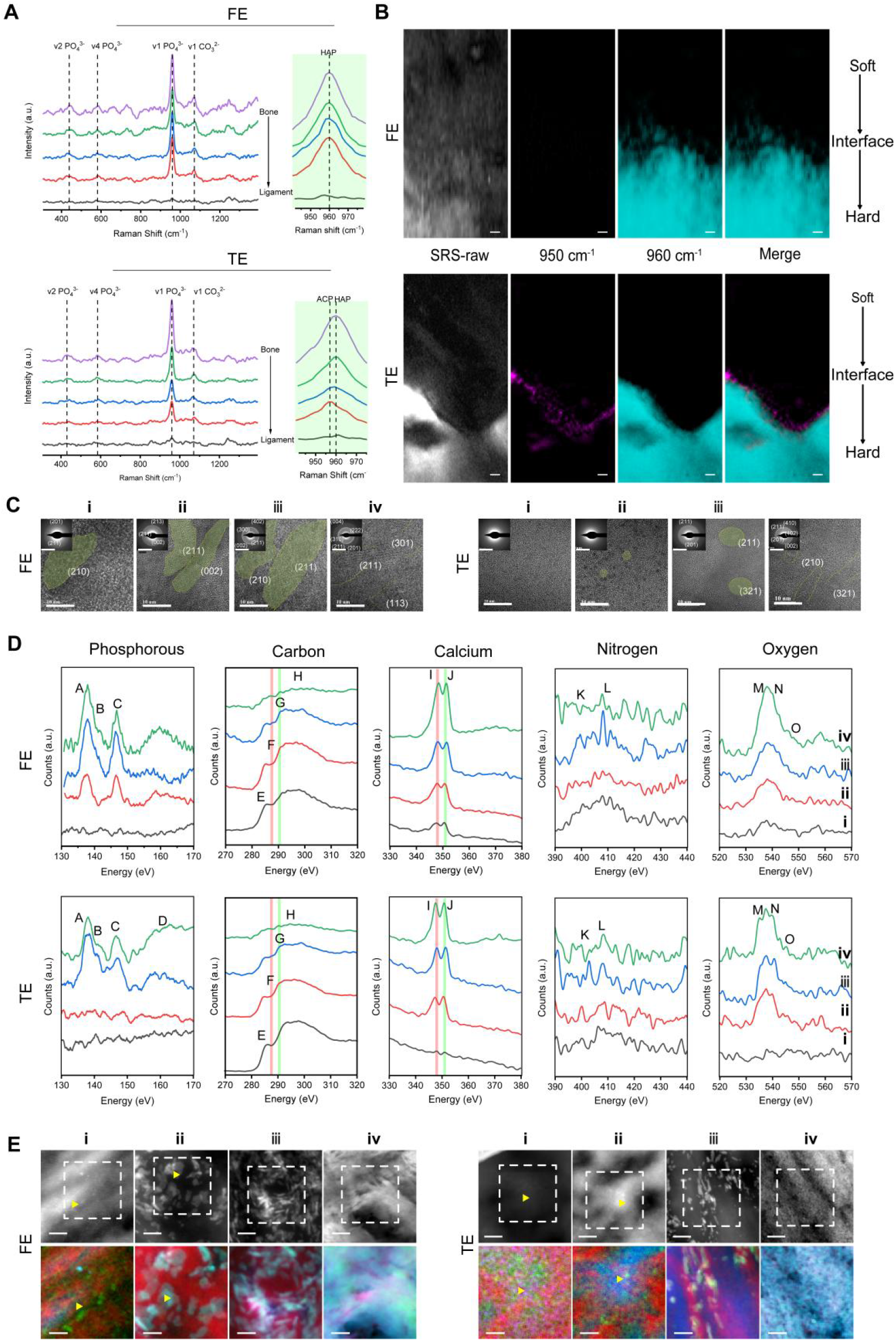
Crystallographic features and chemical analyses of the FE and TE interfaces. (A) Raman spectra of the FE and TE interfaces from ligament to SB. (B) Spectroscopic fingerprint SRS imaging of FE and TE samples showing ACP at 950 cm^−1^ and HAp at 960 cm^−1^ by LASSO unmixing. (C) Corresponding HRTEM and SAED images collected in zone i-iv in Figure 2A revealing mineral assemblies and their crystallinities of FE and TE samples. (D) EELS spectra taken at phosphorus L2,3 edge, carbon K edge, calcium L2,3 edge, nitrogen K edge and oxygen K edge collected from different mineral particles (i–iv) in E. (E) HHADF-STEM images and corresponding composition maps collected at calcium L2,3 edge, nitrogen K edge and oxygen K edge of minerals from different mineral particles (i−iv) in E (red-nitrogen K edge, green-oxygen K edge, blue-calcium L2,3, the original images and processing details see Figure S13). Scale bar in (B), 5 μm; (C, HRTEM images), 10 nm; (C, SAED), 5 1 /nm; (E), 10 nm.

The patterns of mineral assembly at the nanoscale at the FE and TE interfaces were analyzed using high-resolution transmission electron microscopy (HRTEM) and selective electron diffraction (SAED) techniques (Figure 4C). HRTEM images clearly exhibited an increase in the crystallinity of the particles present at both FE and TE interfaces (i-iv). In the FE sample, the crystallization area initially appeared as an elongated strip of 10 nm with poorly crystalline diffraction, eventually fusing. In contrast, in the TE sample, it changed from amorphous, spherical particles with the amorphous scatter of diffuse rings (ACP) to particles with a length of approximately 10 nm ((201) and (211) plane) and then to highly mineralized polycrystalline particles ((002), (102), (201) and (211) plane). Compared to those in the TE sample, the mineral crystals in the FE sample were larger, had a higher aspect ratio, and displayed greater crystallinity. These findings unveiled the nanoscale heterogeneity in the assembly modes of HAp at the FE and TE interfaces, which can enhance force transmission by effectively dissipating energy at the interface.^10^

Further, we used EELS to elucidate the nanoscale chemical environment of various minerals at both FE and TE interfaces (Figure 4D and E; SI Table 1). Our results revealed variations in the content of P (L2,3 edges), C (K edges), Ca (L2,3 edges), N (K edge), and O (K edges) in each particle, consistent with the results obtained through EDX (Figures S12 and S13). It is well known that bone carbonate content is of particular importance in regulating HAp formation, assembly mode, and morphology.^58^ Notably, the C (K-edge) spectra of all particles in the FE sample exhibited four characteristic peaks, while in the TE sample, carbonyl (peak F) mainly appeared in particles i-ii, while particles iii-iv exhibited a higher intensity of carbonate (peak G). EELS maps illustrating the localization of the N, O, and Ca signals are shown in Figures 4E and S13. The nitrogen signal was found to be co-located with the mineral or originates from diffuse “cloudy” structures surrounding the mineral, likely representing the organic ECM component. The coexistence of nitrogen and carbonyl peaks within the granules may be attributed to the presence of proteins during the early stage of mineralization.^59^ Our findings suggest heterogeneity in the essential mineral components of MFC, nay be possibly due to distinct mineralization pathways regulated by organic proteins at the FE and TE interfaces.

### Biomolecular Composition of FE and TE Interfaces

In addition to the transformation of micro/nanostructures and mineral heterogeneity at the nanoscale, the distinctive structural and functional characteristics are defined by variations in biomolecules at the interface from a molecular standpoint. So we employed liquid chromatography-tandem mass spectrometry (LC-MS/MS) to identify the various proteins enriched at the interface (Figure 5A, supplementary file 2). The results revealed that 59 proteins and four proteins exhibited high expression levels in FE compared with the ligament and bone, respectively (Figure 5B and C). In contrast, in tibial samples, 11 proteins and 19 proteins were highly expressed in TE compared to the ligament and bone, respectively (Figure 5D and E). Our findings suggest that the protein expression patterns of FE are more analogous to those of bone tissues, and those of TE tissues are more akin to those of ligament tissues. Nonetheless, these results may have been impacted by incomplete extraction because of the ultra-thin nature of the interface tissue, which may consequently underestimate protein identification.

**Figure 5.**
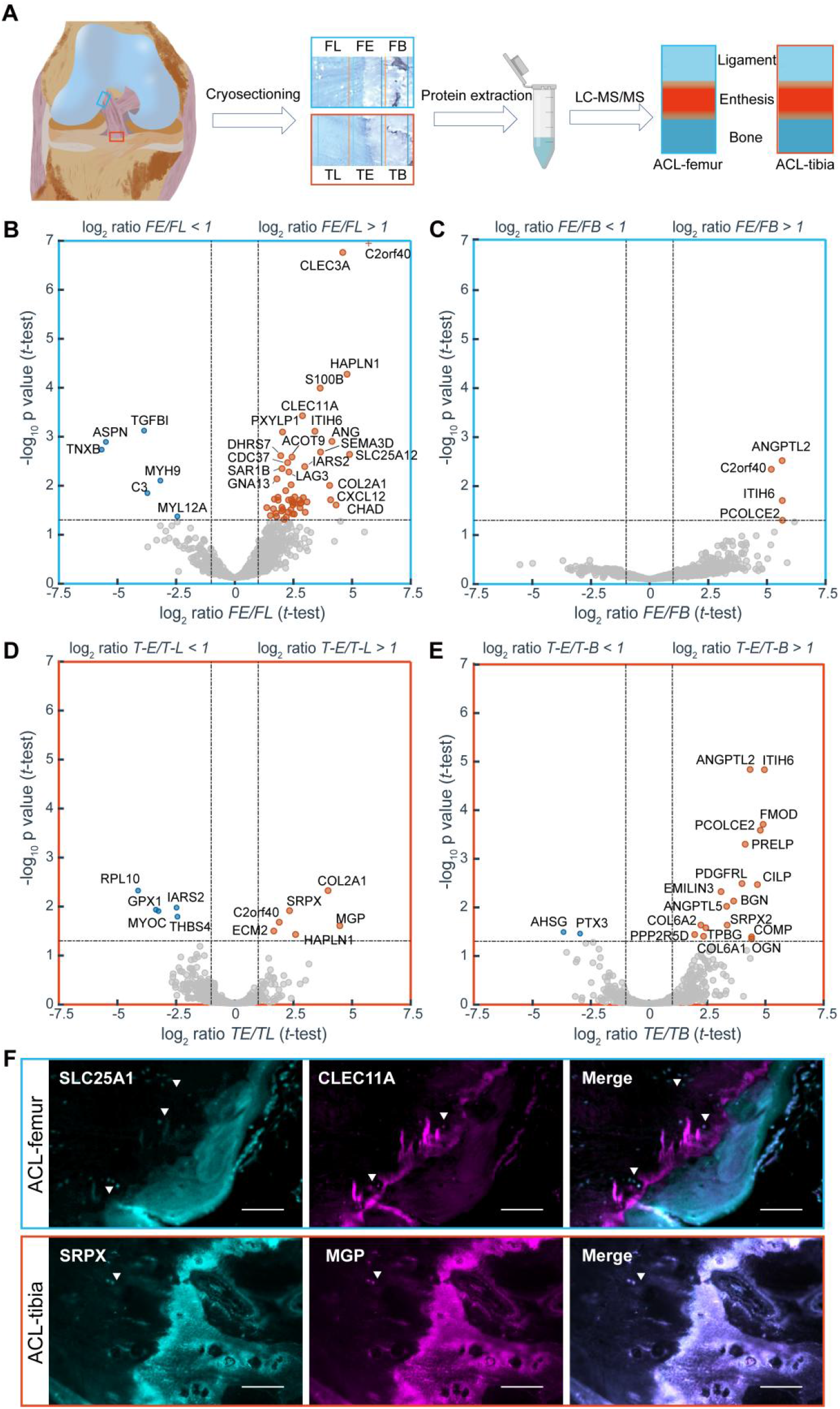
Quantitative proteome analysis of the difference in expression of FE and TE interfaces compared to Ligament and SB. (A) Experimental procedure for the proteomic investigation. Volcano plots of univariate statistical analysis as applied to ligament-bone samples (n = 3 for each tissue). The plots are based on the fold change (log2) and the P-value (-log10) of all proteins identified in (B) proteins from FE and FL; (C) proteins from FE and FB; (D) proteins from TE and TL; (E) proteins from TE and TB. (F) Immunofluorescence staining of SLC24A1 (cyan) and CLEC11A (magenta) in ACL-femur tissues and SRPX (cyan) and MGP (magenta) in ACL-tibia tissues. Scale bar, 20 μm.

Interestingly, in femoral samples, we have identified that the C-type lectin domain containing 11A (CLEC11A) and solute carrier family 25 member 12 (SLC25A12) were significantly upregulated in FE as compared to ligament tissue (Figures 5B and S14). Furthermore, in tibial samples, our analysis revealed that matrix Gla protein (MGP) and sushi repeat-containing protein X-linked (SRPX) were significantly upregulated in TE as compared to ligament tissue (Figures 5D and S14). We have further validated these observations through immunofluorescence staining (Figure 5F). Collagen type II exhibited high expression levels on both interfaces in comparison to the ligament tissue, corroborating the findings of previous studies.^11,60^ Previous studies have suggested that CLEC11A may play a role in promoting osteogenesis by stimulating the differentiation of MSCs into mature osteoblasts,^61^ while MGP may serve as a regulatory protein involved in ACP stabilization and HAP crystallization at the interface.^62^ This may explain the reduced crystallinity observed in TE samples compared to FE samples. Additionally, SLC25A12, a mitochondrial protein, may enhance glycolytic efficiency and muscle fatigue resistance,^63,64^ while SRPX, an ECM protein, may be involved in muscle regeneration and fibrosis,^65–67^ providing a possible explanation for the favorable mechanical response observed at the interface. It should be noted, however, that a detailed understanding of the local distribution of these molecules and their molecular interactions with the encompassing collagen matrix and minerals remains to be explored. Further studies, potentially utilizing gene knockout mouse models, may offer significant elucidation regarding the fundamental influence of these substances on the biomechanical aspects of the ligament-bone interface.

The intricate biological characteristics of the enthesis have captivated the attention of orthopedics and tissue engineers for numerous years.^68^ Recent study in a mouse model has demonstrated the establishment of a mineral gradient at the supraspinatus tendon-to-bone insertion shortly after birth, with an increase in the carbonate content of the apatite mineral at the enthesis over time.^22^ The mineralization of collagen fibrils with carbonated hydroxyapatite leads to the contraction of collagen fibrils, generating substantial stresses of several megapascals and the magnitude of these stresses relies on the type and quantity of mineral present.^69^ These discoveries suggest that the mineralized matrix is crucial for the functionality of the interface. The graded mineralization identified at FE and TE functions to minimize the creation of stress concentrations and enhance load transfer between soft tissue and bone. This facilitates the secure attachment of mechanically dissimilar tissues.^16^ In particular, the development of mature HAps with a fibrous arrangement may enhance the ability of the femoral enthesis to resist shearing forces, while the conversion from ACP to maturing HAps may confer tensile resistance to the tibial enthesis (Figures 2-4). It is widely recognized that mechanical loading plays a vital role in enthesis development and affects the crystallinity within the MFC region of the enthesis.^70,71^ Thus, the variance in mineral content and crystallinity between FE and TE may indicate inherent differences in the loads experienced by these tissue regions. The obtained mineral gradients and the presence of a mixture of crystalline and amorphous calcium phosphate across the interfaces suggest that the observed mineralization is regulated by a complex cellular process involving the release of mineralization precursors or inhibitors from tenocytes and hypertrophic chondrocytes.^72,73^ Future research is needed to investigate both the cellular process and mechanical stimuli responsible for maintaining the nanoscale gradient patterns of calcium phosphate.

### Summary

We conducted a comprehensive investigation into the graded mineralized areas of two highly compliant soft-hard interface structures at FE and TE (Figure 6). Our findings illustrate that the soft-hard transition zone has a width of approximately 20 μm at FE and 10 μm at TE, with both exhibiting an interlocking structure. In both FE and TE, the gradient mineralization zone exhibits an exponential increase in modulus. FEA indicates that FE is favorable for shear, while TE is advantageous for stretching. The distinct mechanical properties of these two interfaces are attributed to the intricate organic and inorganic assembly and the distribution pattern of tissue modulus at multiple scales. FE gradient mineralization is constituted of HAp with varying degrees of crystallinity, whereas TE indicates the transformation from ACP to HAp. The mineral assembly at the FE interface demonstrates high aspect ratio nano-strip HAp in the direction of collagen fiber orientation. The mineral assembly at the TE interface, on the other hand, exhibits a disordered spherical particle gradient distribution with a small aspect ratio. LC-MS/MS analysis confirms the correlation between mineral assemblage and characteristic protein expression at both interfaces. To summarize, the combination of microstructure, micromechanics, and organic and inorganic assembly characteristics elucidates the mechanical advantages and associated mechanisms of FE and TE interfaces. Therefore, this study offers a novel approach to the development of biocomposite interface materials, presenting promising opportunities for ACL reconstruction/repair.

**Figure 6.**
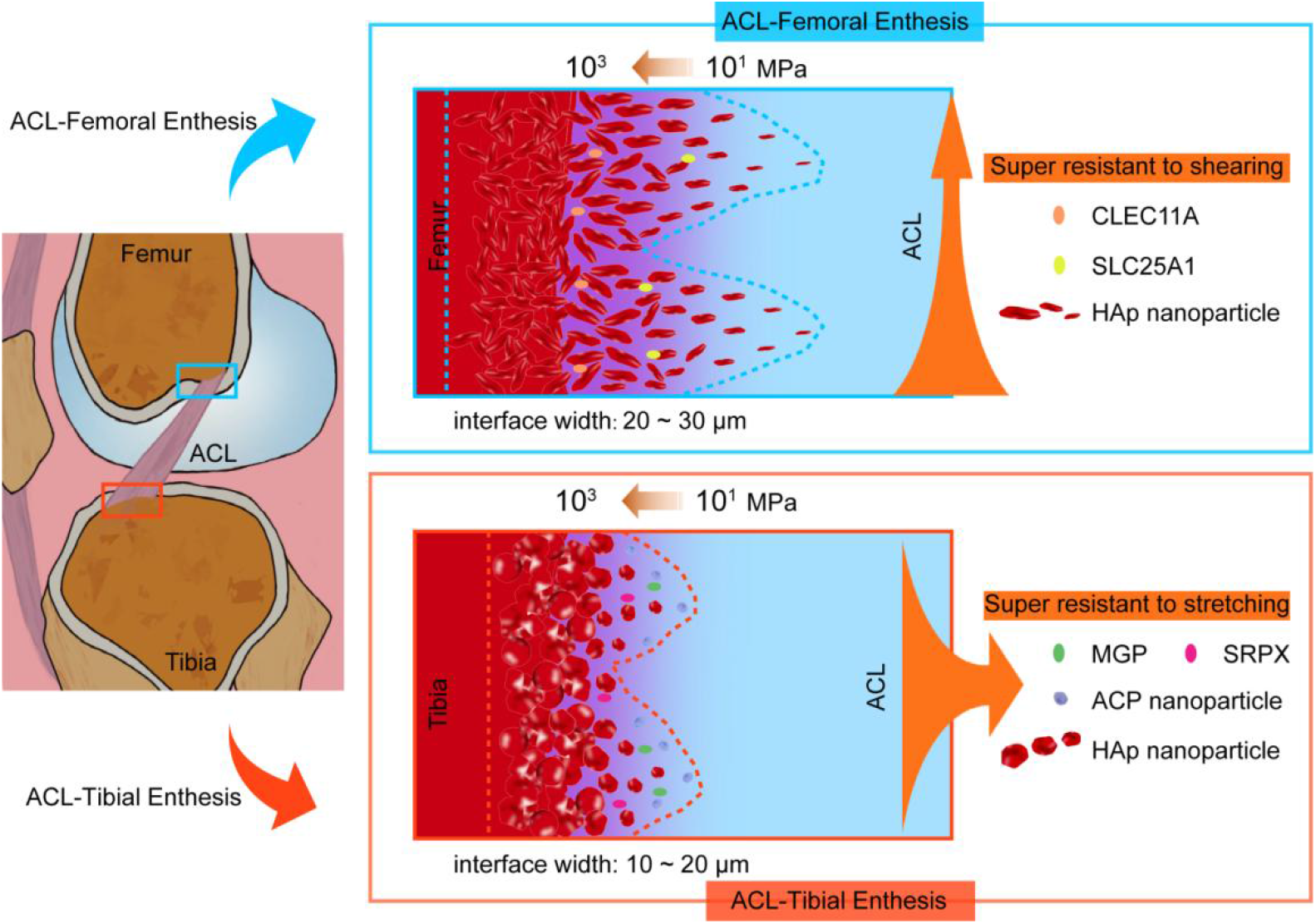
Schematic diagram showing the specific ultrathin interfaces at the FE and TE of human knee joints.

## Materials and Methods

### Sample Preparation

Samples of the knee’s normal ACL-femur and ACL-tibia interface were collected during surgical amputation following the approval from the Ethics Committee of the Second Affiliated Hospital of Zhejiang University School of Medicine. The specimens were obtained from 3 males and 1 female (aged 34, 37, 38, and 39 respectively) with ACL and without macroscopic degenerative changes. The ACL femoral and tibial attachments were carefully collected, which contained ligament, fibrocartilage, and subchondral bone. The tissue was cut into small pieces and later washed with sterile PBS before being stored at −80 ℃ without any further handling or examination. After collection, the samples were split into three sections: one for histology, one for micro-CT and the other for serial sections with different thicknesses for high-resolution characterization (SEM, TEM, FIB-SEM, Raman, etc.) and proteomics.

### Micro-Computed Tomography

To evaluate the microarchitecture of mineralized fibrocartilage, micro-CT (micro-CT, Bruker, Skyscan 1272) was employed. Using a scalpel, the frozen samples were sliced into approximately 3 mm cubes and then immersed in a 2.5% glutaraldehyde solution for 48 hours with continuous stirring. Following the washing process, the samples were submerged in a contrasting agent (Imeron 300). Subsequently, they were subjected to vacuum pressure (1000 mbar) for a duration of 2 minutes to improve tissue penetration. Afterward, the samples were left to incubate in the identical solution for the duration of the night. The samples were rinsed multiple times with 75% ethanol and then placed in a solution of 100% ethanol for subsequent measurement. Subsequently, micro-CT was conducted. Approximately 1200 projections were gathered, each having a pixel dimension of 3 micrometers. Amira 2020 (Thermo Fisher) was employed for the purpose of 3D visualization.

### Histology

Samples of Normal ACL femoral enthesis (FE) and ACL tibial enthesis (TE) were fixed in a 4% paraformaldehyde solution for 48 hours. Subsequently, decalcification was carried out utilizing a 10% EDTA-2Na (w/v) solution over a duration of 4 weeks. Afterwards, both specimens were dehydrated and then encased in paraffin. To identify the ligament-bone interface, measuring 7 μm in thickness were created for Hematoxylin-eosin (HE) and Safranin O (SO) staining.

### Immunofluorescent Staining

Immunofluorescent analysis was performed according to a standard method. Sections with a thickness of 7 μm thick were initially thawed at ambient temperature and rinsed with a PBS solution having a pH of 7.2. Following an overnight thermal antigen retrieval process utilizing a sodium citrate solution (10 mM) at 65°C, the samples were sealed with 5% (wt/v) bovine serum albumin (BSA, Sangon Biotech, China) at ambient temperature for 60 minutes. Next, the FE samples were subjected to incubation with various primary antibodies, including SLC25A1 antibody (1:50, 15235-1-AP, Proteintech) and CLEC11A antibody (1:100, 55019-1-AP, Proteintech). In the same way, the TE samples were left to incubate overnight at 4°C with various primary antibodies, including MGP antibody (1 400, 10734-1-AP, Proteintech) and SRPX antibody (1 200, 11845-1-AP, Proteintech). Following PBS Tween (0.01% Tween) wash, the samples were left to incubated at room temperature for 90 minutes with the corresponding secondary antibodies (diluted 1:200 in PBS): Alexa Fluor® 488 Goat Anti-Rabbit IgG Antibody (Product #A-11008, Invitrogen) and Anti-Mouse IgG Secondary Antibody, Alexa Fluor 546 conjugate (Product #A10036, Invitrogen). Following the rinsing process with PBS solution, the samples were mounted using a fluorescent mounting medium in order to facilitate imaging with a Zeiss 880 laser confocal microscope. Image J was used to analyze the obtained images.

### SEM and EDX Analyses

Fresh FE and TE samples were thawed at ambient temperature, then immersed in a solution containing 2.5% glutaraldehyde, followed by rinsing with a PBS solution. Subsequently, the specimens were cryo-sectioned into slices that were 30 μm thick along the longitudinal orientation of the interfaces. After cryo-sectioning, the slices were washed with DI water to eliminate the OCT, followed by a series of ethanol solutions (20%, 30%, 40%, 50%, 70%, 80%, 90%, 95%, 100%, and 100% (v/v)) for 20 minutes each to facilitate dehydration, and finally left to dry in the air. The samples underwent gold-sputtering and were imaged (Zeiss G300). In order to examine the microstructure of MFC, secondary electrons were collected using a 5 kV accelerating voltage. Acquiring the DDC-SEM images involved the utilization of a 10 kV acceleration voltage. The equipment was equipped with a sensor inside the lens and another sensor that captured secondary electrons and backscattered electrons, respectively. Both modes were used to capture the same area, resulting in the acquisition of DDC-SEM images. The green channel was assigned to the in-lens images, while the red channel was assigned to the backscatter images using Image J. Afterward, the two images were piled on top of each other. Additionally, EDX spectra were obtained in the identical areas utilizing both line and mapping modes to conduct further analysis on the elemental compositions.

### FIB-SEM

Specimen preparation: Using a scalpel, small slices of FE and TE samples were carefully dissected, which contained the regions between nonmineralized to mineralized regions at the ligament-bone interface. Next, the specimens were preserved in a solution of 2.5% glutaraldehyde in PBS for the entire night, then stained with a PBS solution containing 2% osmium tetroxide for a duration of 2 hours. Following multiple rinses with PBS solution, the specimens underwent dehydration using a sequence of acetone concentrations (30%, 50%, 70%, 90%, 100%, and 100%) with each step lasting one hour, except for the 70% concentration which lasted overnight. Ultimately, the specimens were encased in spur resin. To reveal longitudinal sections of the ligament-bone interfaces, all embedded samples were grounded and then polished using a diamond suspension to enhance surface smoothness for the following FIB-SEM serial imaging.

Data collection: The Leica EM trimmer was used to carefully trim resin blocks until the black tissue surface in the block became visible. In order to visualize the region of interest, a scanning electron microscope (Thermo Fisher, Teneo VS) equipped with an ultramicrotome within its specimen chamber was employed. This enabled us to precisely trim the resin blocks and simultaneously capture the electron microscopic image of the sample. Once the region of interest was identified, imaging was conducted utilizing dual-beam scanning electron microscopy (Thermo Fisher, FIB Helios G3 UC). The data-gathering process was conducted using the sequential-surface perspective method, with a slice thickness of 6 nm at 30 keV and 0.43 nA current. Afterwards, every consecutive facial image was captured in backscatter mode (BSE) utilizing an ICD detector, with a 2 kV acceleration voltage and a current of 0.2 nA. The resolution of the image store was set to 3072×2048 pixels, with a dwell time of 15 μs and a pixel size of 4.25 nm. ImageJ was used to measure the aspect ratios.

Segmentation and processing of images: Amira 2020 (Thermo Fisher) was utilized to align and crop the image stack. The manual process of segmenting and labeling the cuticle and sensilla was performed, and surface-generation tools were employed to calculate the surfaces of structures of interest.

### Raman Spectroscopy

Raman spectroscopy with a confocal Raman microscope (LabRAM HR Evolution, Horiba Co., Ltd.) equipped with a 532 nm laser was utilized to measure the chemical components across FE and TE samples. An EMCCD detector with a spectral resolution of approximately 1 cm^−1^ was utilized to collect spectra within the range of 200 to 1900 cm^−1^. For the point mode, the spectra were acquired with a 5 s accumulation time. In the case of Raman mapping, a continuous scan was conducted with a spatial resolution of approximately 1 μm and an accumulation time ranging from 0.1 to 0.5 s. There was no observed deterioration of the sample when using this parameter.

### Stimulated Raman Scattering Microscopy

Stimulated Raman scattering microscopy was performed to detect and visualize the amorphous calcium phosphate (ACP) and hydroxyapatite (HAp) across FE and TE samples using a Multimodal Nonlinear Optical Microscopy System (UltraView, Zhendian (Suzhou) Medical Technology Co., Ltd). A hyperspectral scan step size of 0.5 cm^−1^ was used to collect spectra within the range of 920∼1002 cm^−1^. For imaging, 400 × 400 pixels (0.52 μm/pixel) are acquired with a 30-μs dwell time. The least absolute shrinkage and selection operator (LASSO) method was utilized for spectroscopic fingerprint SRS imaging.^1^ Each image represents substances of different spectra.

### TEM, EDX, HAADF-STEM, EELS and SAED Analyses

The samples were immersed in a 2.5% glutaraldehyde solution for 24 hours, rinsed with a PBS solution, treated with 1% OsO_4_ stain, dehydrated, and subsequently enclosed in a spur resin. Afterwards, the samples were sliced into thin sections using a Lerca EM UC7 ultratome with a thickness of 100 nm. These sections were then positioned onto copper grids with a mesh size of 200 and analyzed using a Super-X EDX detector-equipped aberration-corrected scanning transmission electron microscope (FEI Titan G2 80-200 microscope). The examination was conducted at a voltage of 200 kV. Regions of interest were examined using HAADF-STEM, TEM-EDX, and SAED patterns. Images of EELS spectra were collected in the range of 100-600 eV to investigate the characteristic edges of the elements of interest (P, C, Ca, N, O). Principal component analysis was applied to calibrate, normalize, subtract background, and process each spectrum. Prior to conducting each experiment, the electron probe of the microscope was calibrated with a DCOR plus spherical aberration corrector by utilizing a gold standard sample. No apparent harm was detected during the experiments. The HAADF-STEM image, SAED, and EELS analysis were conducted using Gatan Digital Micrograph software.

### Proteomics

Sample preparation: Cryo-sectioning was used to prepare ligament, interface and SB tissues from FE and TE samples. It should be mentioned that the difference between interface, ligament and SB were mostly larger than our results, as the interface samples were not completely differentiated from ligament and SB due to the absence of labeling. In the tube, a total of six types of tissue were gathered and then sliced into small fragments. After rinsing with PBS, the tissue samples were scraped off the slides and transferred into 0.6 ml centrifuge tubes. Subsequently, 20 μl of enzyme digestion buffer (0.1M ammonium bicarbonate) was added to each tube. The mixture was then heated at 95℃ for 10 minutes, followed by the addition of 1 μl of trypsin (1 μg/μl). After shaking and mixing, the samples were incubated at 37℃ overnight (> 12 h). Next, they were subjected to centrifugation at 14000 g for 15 minutes and the resultant supernatant was gathered. The supernatant was then desalted and utilized for peptide concentration determination using a Nanodrop spectrophotometer.

LC-MS/MS analysis: The peptides were dissolved in the liquid chromatography mobile phase A (0.1% (v/v) aqueous solution of formic acid) and subsequently separated utilizing an UltiMate 3000 nanoLC ultra-high performance liquid chromatography apparatus. The self-filling column used in chromatography had dimensions of 25 cm length and 100 μm inner diameter, packed with C18 material of 1.9 μm. The first mobile phase, referred to as A, consisted of an aqueous solution containing 0.1% formic acid. The second mobile phase, referred to as B, consisted of an aqueous solution containing 0.1% formic acid and 98% acetonitrile. Liquid phase gradient settings: 0-3 min, 3%∼10% B; 3-40 min, 10%∼24% B; 40-52 min, 24%∼38% B; 52-56 min, 38%∼80% B; 56-60 min, 80% B. The rate of flow was kept constant at 450 nL/min.

After being separated by the UHPLC system, the peptides underwent ionization in the NSI ion source and were subsequently analyzed using the Orbitrap Exploris 480 mass spectrometer. A voltage of 2.0 kV was applied to the ion source, and the high-resolution Orbitrap was utilized to detect and analyze both the peptide parent ions and their secondary fragments. The initial range for mass spectrometry scanning was established as 400-1200 m/z with a scan resolution of 60,000. Additionally, the Orbitrap scan resolution was set at 15,000. The data acquisition mode employed a data-dependent scanning (DDA) program with a cycle time of 1 s. To fragment the HCD collision cell, a fragmentation energy of 27% was utilized. In order to enhance the efficient utilization of the mass spectrum, the automatic gain control (AGC) was adjusted to 100%, the signal threshold was established at 50,000 ions/s, the maximum injection time was limited to 22 ms, and a dynamic exclusion time of 30 s was implemented for the tandem mass spectrometry scan to prevent redundant scans of the parent ions. FAIMS Pro’s compensation voltage was adjusted to −45 V.

Database search: MaxQuant (v1.6.15.0) was utilized to search the data obtained from secondary mass spectrometry. The database was searched using UniProt Human (20387 sequences). To calculate the false positive rate (FDR) caused by random matching, the inverse library was included. Additionally, the common contamination library was added to eliminate the impact of contaminated proteins on the identification results. The enzyme cut mode was set to Trypsin/P, with a maximum of 2 missed cut sites. The mass error tolerance for the primary parent ion was set to 20 ppm for the First search and 5 ppm for the Main search. The mass error tolerance for the secondary fragment ion was set to 0.02 Da. Fixed modification was assigned to cysteine alkylation, while oxidation of methionine and acetylation of protein N-terminal were considered as variable modifications. The FDR for protein identification and peptide-spectrum match (PSM) identification was established at 1%.

Protemoics data analysis: The limma R package was used to identify the proteins that showed differential expression, using PSMs information as input. The data underwent log2-transformation and was normalized using the normalize Between Arrays function. Next, the lmFit function was employed to establish a linear model for every protein. For every protein, the moderated t-statistic of eBayes was computed, and the p-values were adjusted for multiple testing using the positive FDR through the contrasts fit and eBayes functions. Proteins exhibiting differential expression were characterized by a fold change greater than 2 and an adjusted p value below 0.05.

### Nanoindentation

Fresh FE and TE samples were dissected into small pieces about 200 μm-thick along the longitudinal direction of the interfaces and immersed in a PBS solution for further measurement. Nano-indentation tests were performed using a nano-indenter (PIUMA, Optics 11, Amsterdam Netherlands) to investigate indentation behavior. The soft cantilever (radius: 10 μm, stiffness: 159 N/m) was utilized to measure the elastic modulus of tissues. A nanoindentation test matrix (step: x and y-2×2 μm) was conducted on the samples to analyze the modulus transition of the ACL-femoral and ACL-tibial interfaces. In order to replicate the physiological conditions, all measurements were conducted in a PBS solution.

### AFM

To characterize the nanomechanical attributes of FE and TE samples, AFM was utilized for conducting Force-Displacement (FD) measurements. 150-um-thick cryo-cut samples were washed and immersed in PBS solution for further evaluation. The Cypher ES AFM (Oxford Instruments Asylum Research, United States) along with Si cantilevers (AC160TS-R3, Olympus, Japan) were employed. With a back angle α of 35°, the spring constant was 26 N/m and the tip radius measured 7 nm. Data was collected for a 40×40 array of FD measurements in three distinct regions within a 5×5 μm^2^ area for both FE and TE samples. The approach speed of the tip was 3 μm/s until a force of 6 μN was attained, after which the tip was retracted at the identical speed. To fit Young’s modulus, the FD data were analyzed assuming a Hertz model with a tapered end.

### Finite Element Analysis

To investigate the role of the spatial arrangement of the modulus and insertion depth at the FE and TE interfaces, we built a finite element model to investigate their advantages.

Geometric modeling: In the part of Abaqus, the stereo model was established according to the dimensions shown below (Figure S7A).

Partition the computational grid: Both parts of the model adopted a hexahedral and tetrahedral mixed mesh, dominated by hexahedrons for mesh partitioning (Figure S7B).

Define material properties: The ligament part was defined as isotropic elastic with Young’s modulus of 30 MPa and a Poisson’s ratio of 0.3. The material properties of the transition zone were defined based on the data curve obtained from nanoindentation experiments; that was, different material properties were assigned to the corresponding elements according to the different central positions of the grid cells (the average coordinate value of the surrounding nodes). The material properties of individual elements were isotropic elastic bodies with a Poisson’s ratio of 0.3 and Young’s modulus varying in the range of 39.72 to 760 MPa for FE and 27 to 1261 MPa for TE, depending on their locations.

Build the assembly and set up interactions: The two parts were assembled according to their relative position, and the interfaces were bonded.

Define the boundary conditions: The surface close to SB was set as the fixed end (Encastre) (Figure S7C).

Apply load: Displacement loads of 0, 30, 60, and 90 degrees were applied to the tendon end (the same percentage, strain = total displacement divided by model original length = 10%, 20%) and submitted for calculation (Figure S7D).

### Statistical Analysis

The data were presented as the mean ± SD. A one-way ANOVA analysis (Tukey’s posthoc test) was conducted to assess the disparities among the values. A significance level of less than 0.05 was deemed statistically significant.

## Author Contributions

The manuscript was written through contributions of all authors. All authors have given approval to the final version of the manuscript.

## Notes

The authors declare no competing financial interest. ACKNOWLEDGMENT We thank Xi Zheng from Zhejiang University for SEM imaging and EDX analysis; we thank Jiansheng Guo from the Center of Cryo-Electron Microscopy, Zhejiang University for FIB-SEM imaging; we thank Shoupu Yi from Zhendian (Suzhou) Medical Technology Co., Ltd for stimulated Raman spectroscopic imaging; and we thank Guoqing Zhu from Zhejiang University for the HRTEM and SAED imaging.

