## Supplementary material for "Distinct Microstructural Heterogeneities Underpin Specific Micromechanical Properties in Human ACL Femoral and Tibial Entheses": Figure S1-14, Table S1

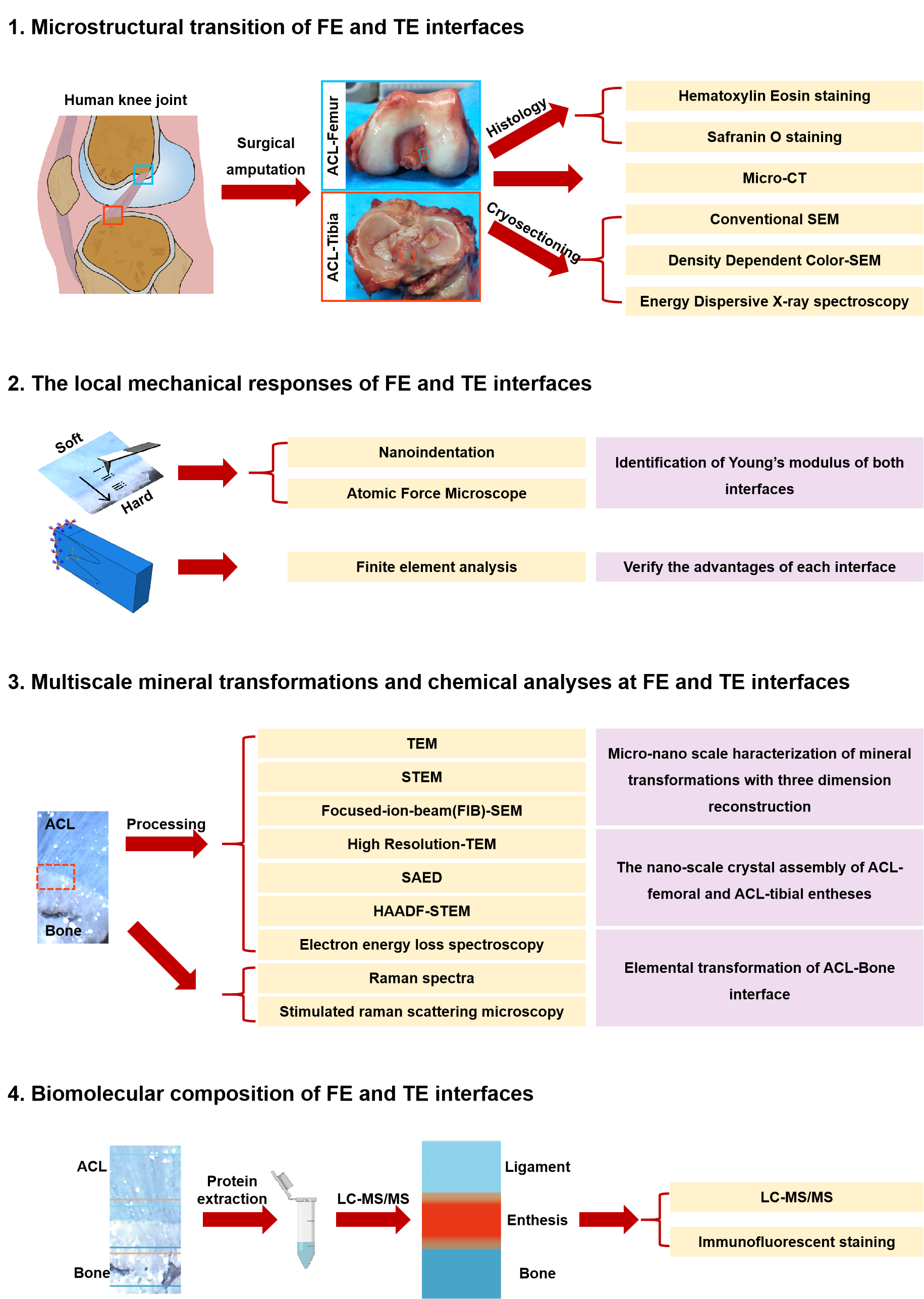

**Figure S1.** Workflow chart of this study. Abbreviations, FE, ACL-femoral enthesis; TE, ACL-tibial enthesis; SAED, selected area electron diffraction. The red rectangle in 3 was the region of interest.

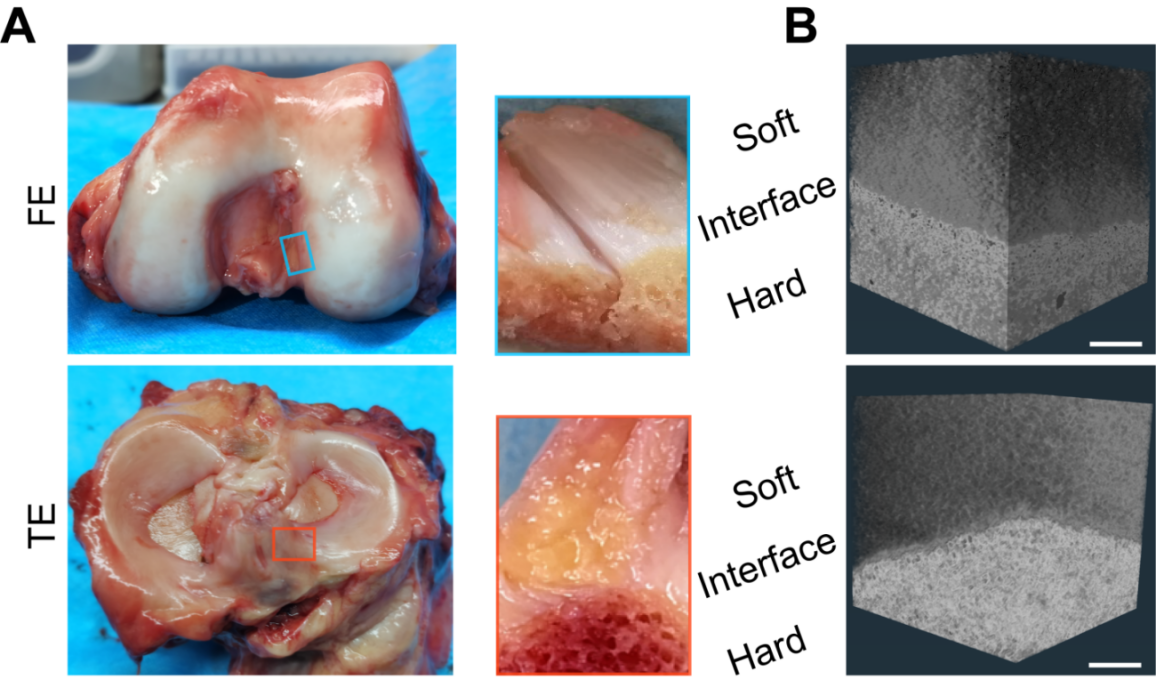

**Figure S2.** (A) Gross appearance of human FE and TE samples. (B) Micro-CT of FE and TE samples. Scale bar in (B), 30 μm

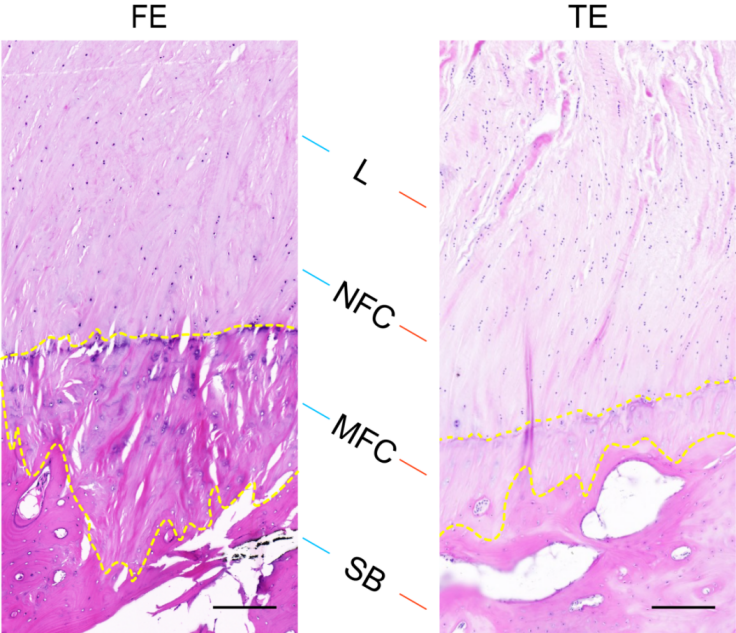

**Figure S3.** HE staining of the FE and TE tissues showing four layers, ligament, NFC, MFC and SB. Scale bar, 200 μm

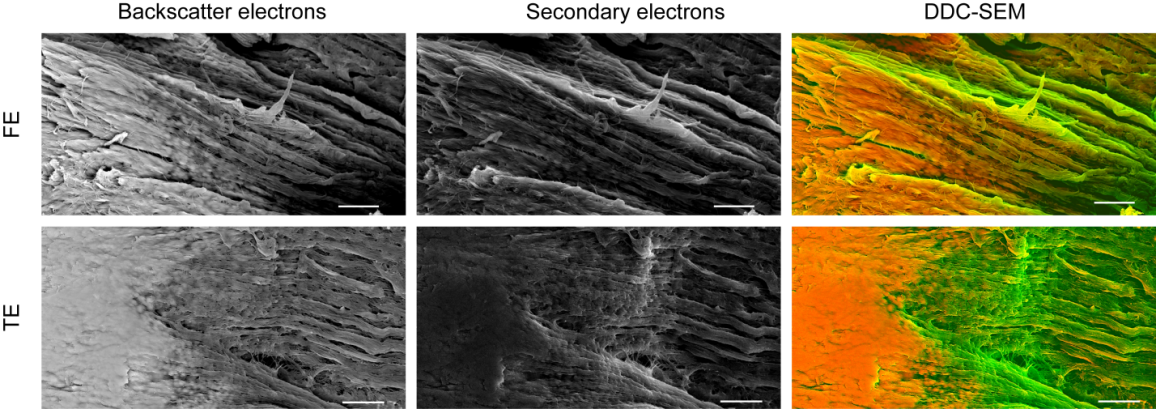

**Figure S4.** Image processing procedures employed to creat a density dependent colour scanning electron micrographs (DDC-SEM) showed in Figure 1D. The backscattered electron images and secondary electron images were assigned to be red and green channels, respectively, followed by stack command to acquire a combined images. After that, the combined images were converted to single RGD images. The final output images clearly presented mineral morphologies and distributions across the interface regions in FE and TE. Scale bar, 5 μm

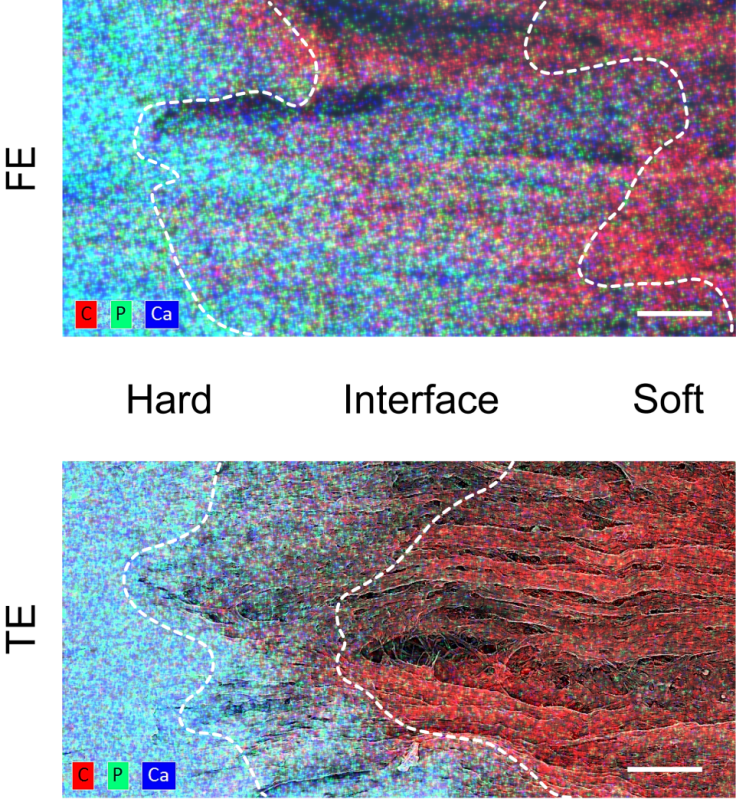

**Figure S5.** The corresponding EDX elemental maps of the regions in Figure 1D and S3. Ca (blue), P (green), and carbon (red) were distributed hierarchically throughout the interface. The Ca/P-rich region was complementary to the C-rich region. Scale bar, 5 μm

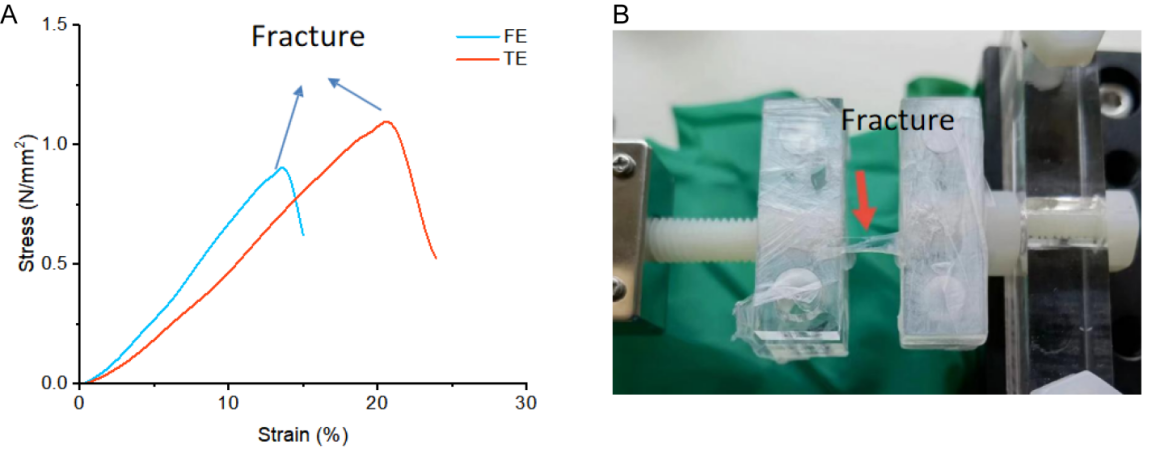

**Figure S6.** (A) Typical specific strength-strain curve of FE and TE sample. (B) Photograph showing the fracture site of FE and TE samples when stretched.

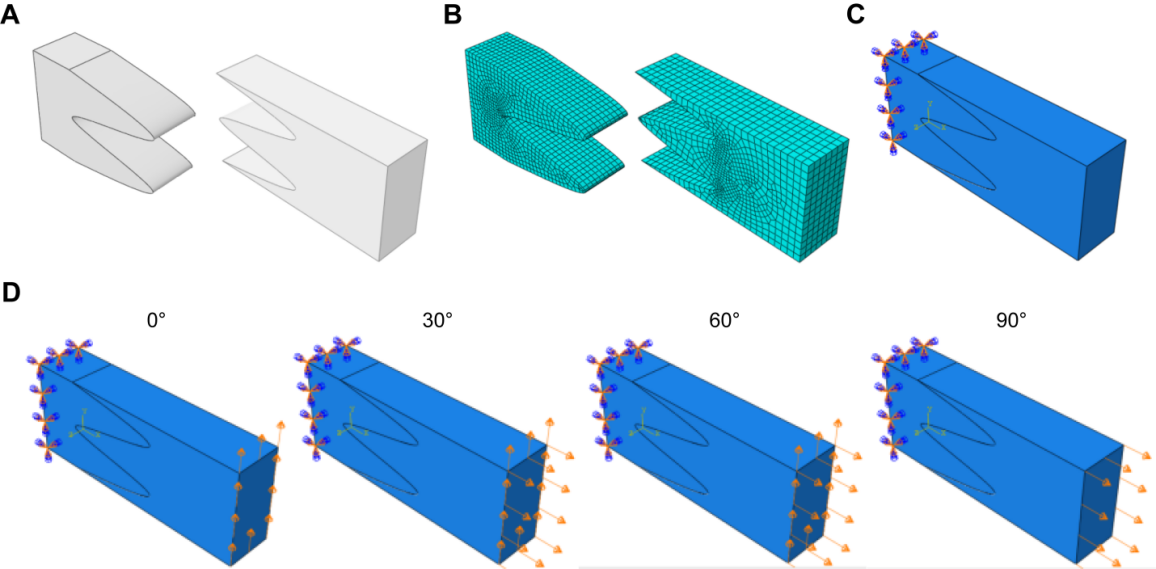

**Figure S7.** Finite element analysis. (A) Geometric modeling, transition zone (left) and tendon (right). (B) Partition the computational grid. (C) Define material properties, build the assembly and set up interactions. The surface close to SB was set as the fixed end. (D) Displacement loads of 0, 30, 60 and 90 degrees were applied to the tendon end (the same percentage, strain = total displacement/model original length = 10%, 20%, 30%) and submitted for calculation.

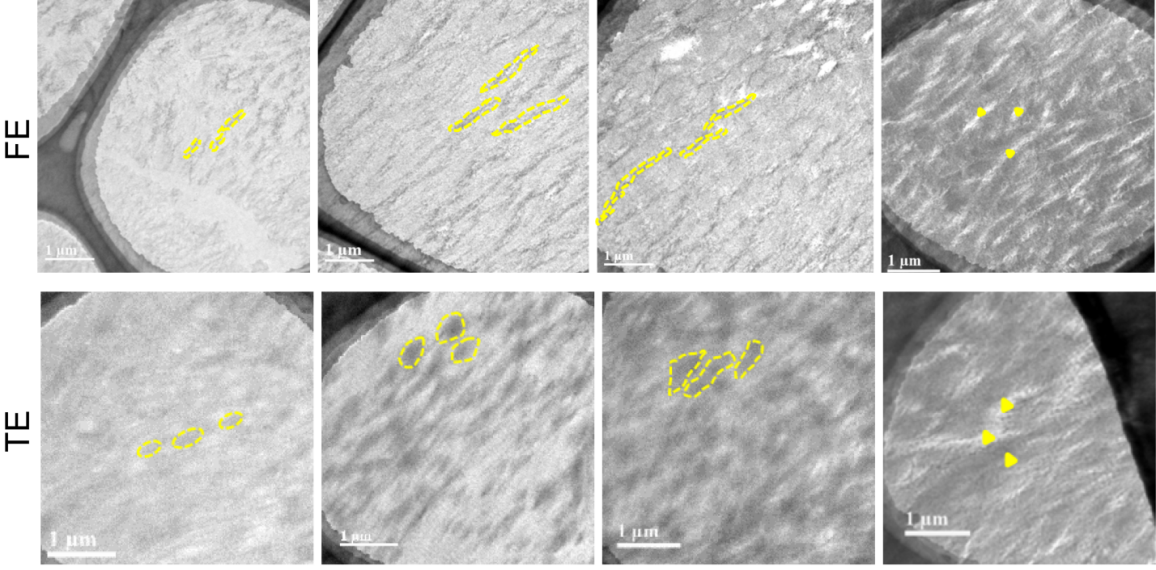

**Figure S8.** TEM of ACL-bone interface tissues presenting gradual mineral distributions with distinct morphologies. Scale bar, 1 μm.

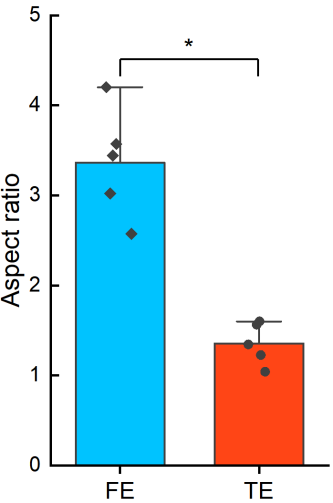

**Figure S9.** Aspect ratios of minerals in Figure 3D (zone i)

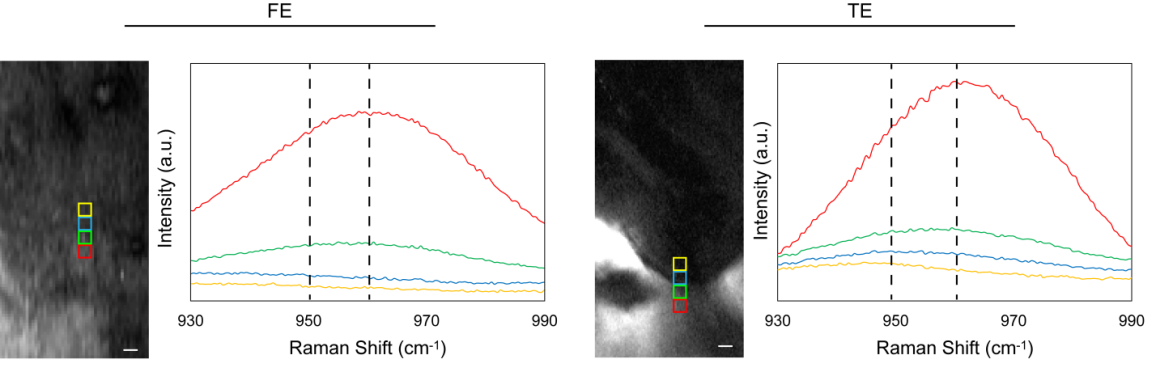

**Figure S10.** SRS spectra of the FE and TE interfaces

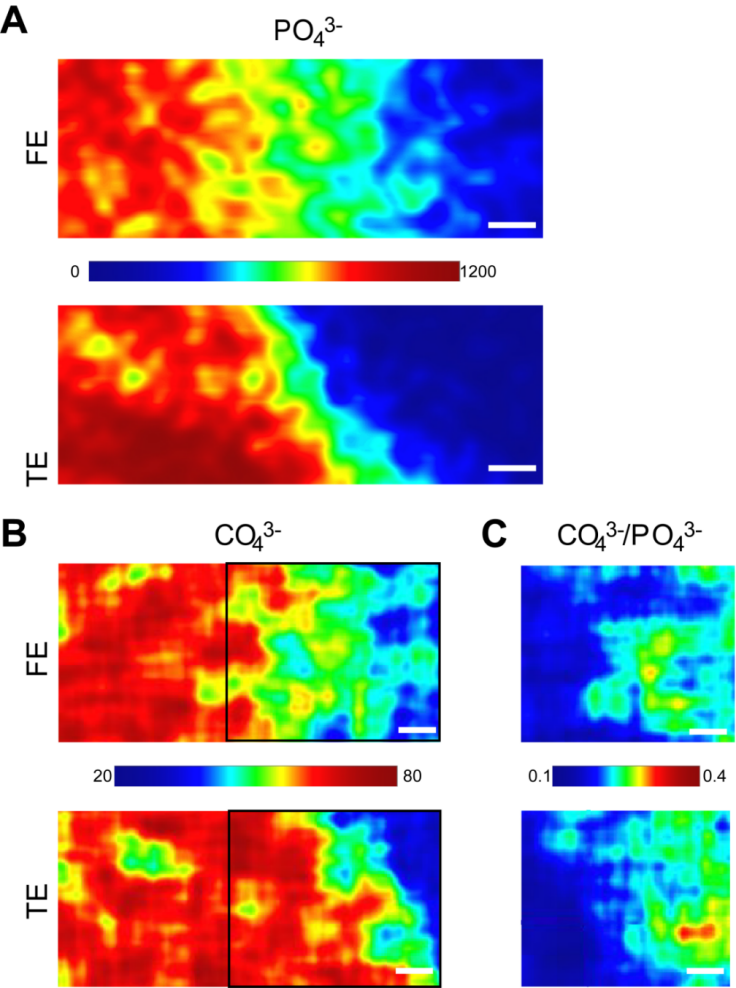

**Figure S11.** Collected Raman maps showing HAp contents (960 cm^−1^, A), CO_3_^2−^ substitution (1071 cm^−1^, B), and CO_3_^2−^/PO_4_^3−^ ratios (C) revealing ionic substitution of minerals across the interface for different samples. Spatial resolution was 1 μm. Scale bar in (A-C), 4 μm.

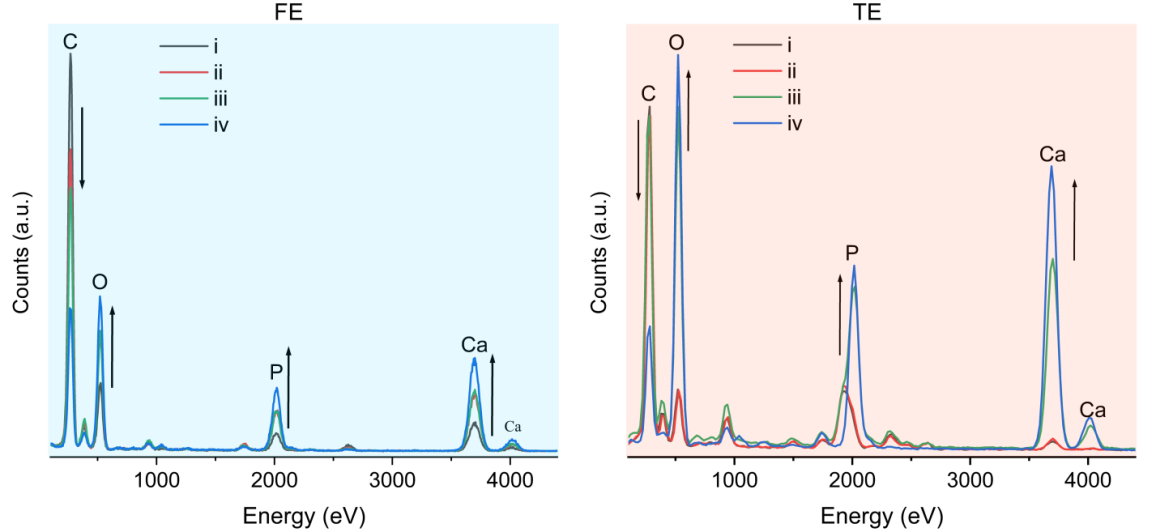

**Figure S12.** TEM-EDX spectra collected from minerals in zone i-iv in Figure 3A showing the elemental changes of FE and TE.

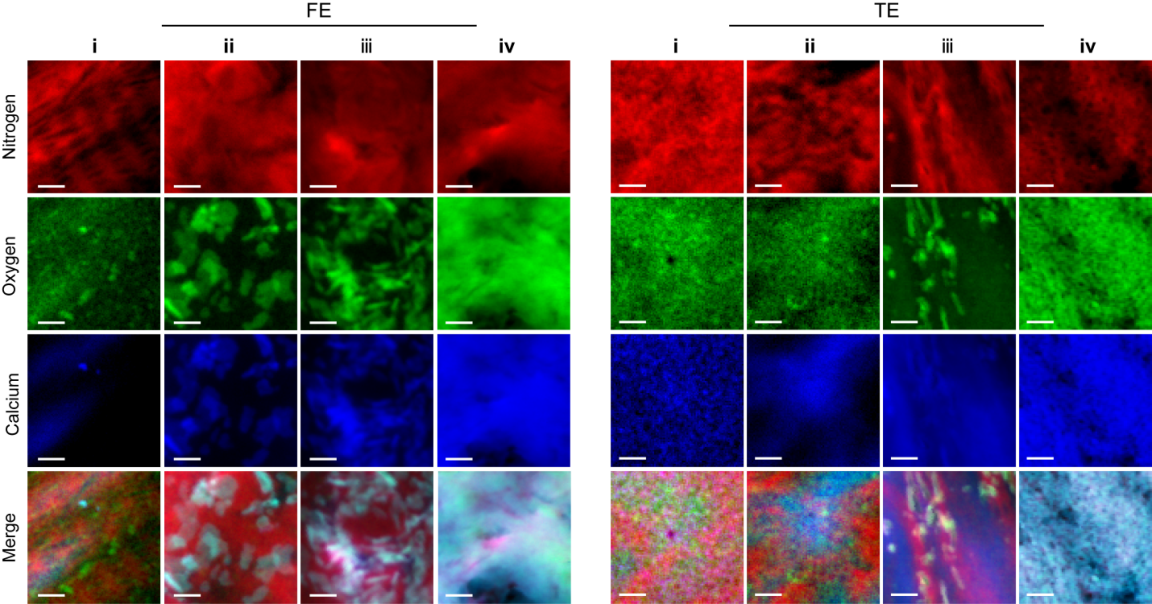

**Figure S13.** EELS mapping of calcium (blue), oxygen (green), and nitrogen (red) of the selected areas in Figure 3G. Scale bar, 10 nm.

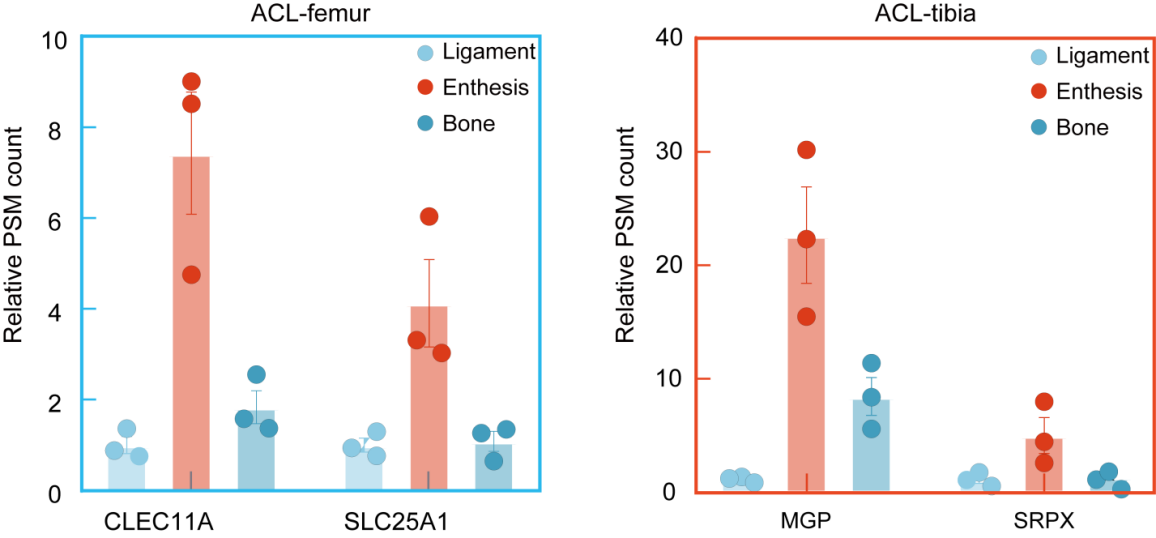

**Figure S14.** Expression of CLEC11A and SLC25A1 in different ACL-femoral tissues, and expression of MGP and SRPX in different ACL-tibial tissures.

**Table S1.** Approximate transitions in phosphorus L_2,3_ edge, carbon K edge, calcium L_2,3_ edge, nitrogen K edge and oxygen K edge fine structure and assignments of peaks.^2^

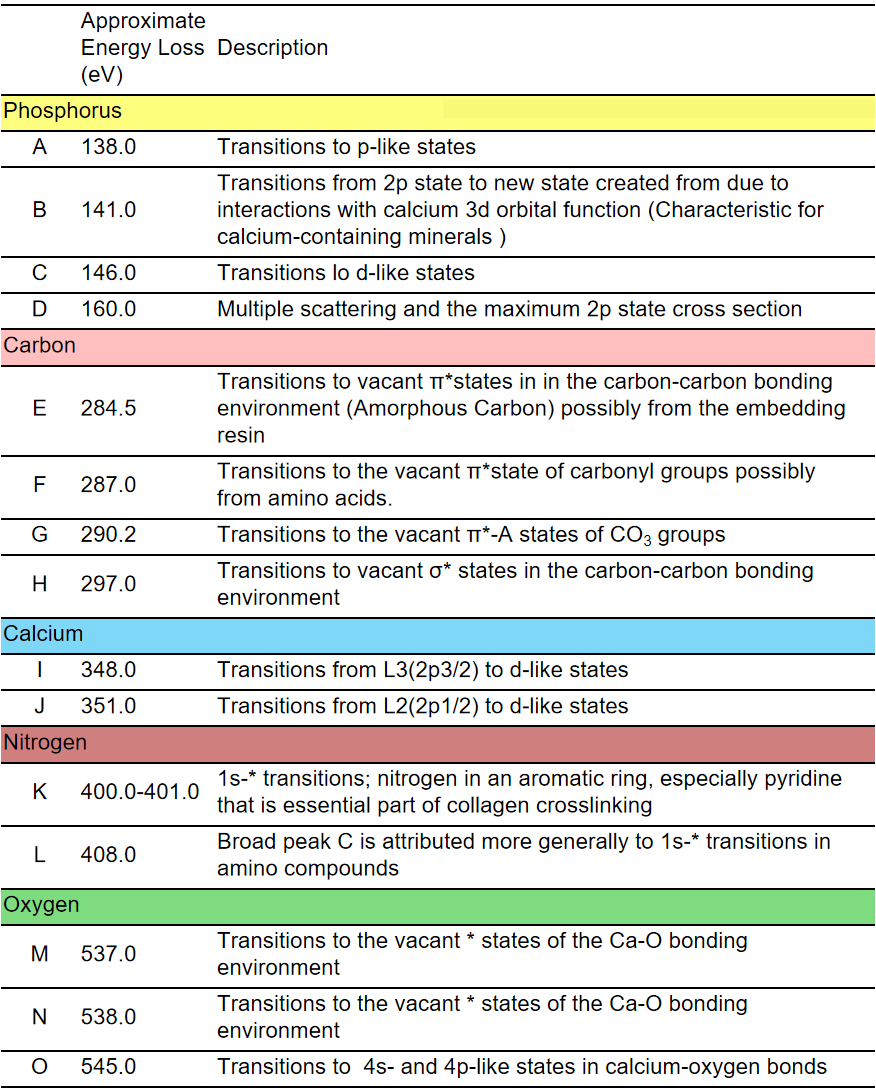

**References**

| (1) | Lin, H.; Lee, H. J.; Tague, N.; Lugagne, J.-B.; Zong, C.; Deng, F.; Shin, J.; Tian, L.; Wong, W.; Dunlop, M. J.; Cheng, J.-X. Microsecond Fingerprint Stimulated Raman Spectroscopic Imaging by Ultrafast Tuning and Spatial-Spectral Learning. Nat Commun 2021, 12 (1), 3052. https://doi.org/10.1038/s41467-021-23202-z. |
| --- | --- |
| (2) | Nitiputri, K.; Ramasse, Q. M.; Autefage, H.; McGilvery, C. M.; Boonrungsiman, S.; Evans, N. D.; Stevens, M. M.; Porter, A. E. Nanoanalytical Electron Microscopy Reveals a Sequential Mineralization Process Involving Carbonate-Containing Amorphous Precursors. AC Nano 2016, 10 (7), 6826–6835. https://doi.org/10.1021/acsnano.6b02443. |
